# Moderate amounts of epistasis are not evolutionarily stable in small populations

**DOI:** 10.1101/752535

**Authors:** Dariya K. Sydykova, Thomas LaBar, Christoph Adami, Claus O. Wilke

## Abstract

High mutation rates select for the evolution of mutational robustness where populations inhabit flat fitness peaks with little epistasis, protecting them from lethal mutagenesis. Recent evidence suggests that a different effect protects small populations from extinction via the accumulation of deleterious mutations. In drift robustness, populations tend to occupy peaks with steep flanks and positive epistasis between mutations. However, it is not known what happens when mutation rates are high and population sizes are small at the same time. Using a simple fitness model with variable epistasis, we show that the equilibrium fitness has a minimum as a function of the parameter that tunes epistasis, implying that this critical point is an unstable fixed point for evolutionary trajectories. In agent-based simulations of evolution at finite mutation rate, we demonstrate that when mutations can change epistasis, trajectories with a subcritical value of epistasis evolve to decrease epistasis, while those with supercritical initial points evolve towards higher epistasis. These two fixed points can be identified with mutational and drift robustness, respectively.

## Introduction

When a population is in mutation–selection balance, it is able to maintain its mean fitness while still generating genetic variation that may increase its fit to the environment via adaptive mutations [1]. However, this balance between the evolutionary forces of selection and mutation can sometimes be precarious. When mutation rates become too high, for example, mutations can overpower selection leading to the extinction of a population via lethal mutagenesis [2]. Similarly, when population size dwindles, selection can become so weak that deleterious mutations cannot be eliminated, leading to fitness decline via Muller’s ratchet [3] or population extinction through a mutational meltdown [4]. Populations can adapt to high mutation rates and/or small population sizes by evolving “mutational robustness” [5] or “drift robustness” [6–8]. Populations evolve mutational robustness by moving onto flat fitness peaks, where they experience a reduction in maximum fitness counterbalanced by an increased fraction of new mutations that are either neutral or have a small fitness effect [9, 10]; this phenomenon is often referred to as the “survival-of-the-flattest” effect [9]. Robustness to drift, on the other hand, appears to involve favoring fitness peaks that have steep flanks, enabled by mutations that are synergistic in their deleterious effect [6, 7], while *reducing* (rather than increasing) the likelihood of mutations with small effect, and increasing the fraction of mutations that are lethal. Interestingly, a recent re-analysis of the survival-of-the-flattest effect has shown that an increase in the fraction of lethal mutations is also seen in the response to high mutation rates [10], suggesting that resistance to drift and resistance to mutations are intertwined (see also [8]).

The threat of high mutation rates and small population sizes to genetic survival is particularly real for populations that periodically undergo bottlenecks during transmission between hosts and cannot rely on sexual recombination to protect against gene loss, such as the mitochondria of the salivarian Trypanosomes *T. brucei* and *T. vivax* [11]. For those organisms, population size often drops into the single digits [12] while mutation rates are elevated due to oxidative stress [13]. High mutation rates and small population sizes are also important for viral populations. Mutational robustness (and possibly drift robustness) has been observed in some strains of the RNA virus vesicular stomatitis virus (VSV) that differ in the rate at which deleterious mutations accumulate at small population size [14].

How genomes respond to mutations is determined to a large extent by how mutations interact. In general, the effect of a mutation on host fitness is influenced by the genetic background within which that mutation occurs, a phenomenon known as epistasis [15]. Epistasis has a *direction*: the effect of a pair of mutations can either be larger or smaller than what is expected from a single mutation, so that the deleterious effect of two mutations can be either amplified (synergistic), or buffered (antagonistic). The average direction between pairs (also-called directional epistasis, see for example [16]) plays an important role in determining linkage equilibria [17, 18], canalization [19, 20], as well as theoretical investigations of the origin of sex [21–23]. Epistasis has been measured quantitatively for a number of model organisms, and both antagonistic and synergistic trends have been observed [24–30].

When faced with changed conditions, one of the ways in which populations can adapt is by changing the way information is encoded in the genome, leading to changes in epistasis [31]. Here we study the impact of epistasis on both drift and mutational robustness in a simple model fitness landscape. We show that whether a population predominantly displays drift or mutational robustness is largely determined by the average value of directional epistasis: populations occupying a peak with synergistic epistasis above a critical value will tend to evolve towards drift-robust peaks (by moving towards peaks with increased positive epistasis), while those inhabiting peaks with sub-critical epistasis will respond by lowering epistasis until mutations are mostly neutral, consistent with mutational robustness. Thus, evolutionary trajectories for populations under evolutionary stress will bifurcate towards drift-robust or mutationally-robust fixed points.

## Model

We study a simple fitness landscape in which the wild type genotype resides on a fitness peak with a height of 1, and the fitness of a *k*-mutant is given by

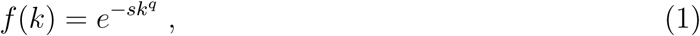

where *s* = −log *f*(1) is the mean effect of a deleterious mutation to the wild type, *q* determines the degree of directional epistasis, and genotypes have a finite number of binary loci *L* (see, e.g., [16])^1^. In such a model *q* = 1 signals absence of epistasis (i.e., the fitness landscape is multiplicative), *q >* 1 describes a peak with synergy between deleterious mutations (synergistic epistasis), and *q <* 1 is indicative of buffering mutations (antagonistic epistasis). When *q >* 1 we sometimes speak of *negative* epistasis (because the combined two-mutant fitness is lower than the multiplicative expectation), while *q <* 1 indicates *positive* epistasis (the double-mutant is higher in fitness than expected on the basis of the single-mutation effect). Of course, a model that only treats the mean epistatic effect between mutations using a single parameter *q* has significant limitations. In particular, such a model cannot capture effects that are due to a distribution of pair-wise epistatic effects (something that can be delivered by an NK model, for example [33]). Furthermore, we ignore here all the subtleties of sexual reproduction, which can also affect how epistasis evolves. The loss of realism is offset by our ability to control the parameters of such an effective model precisely (*s* and *q*), which in more sophisticated models depend on each other. Furthermore, an analysis in terms of asexual processes is warranted for those genomic stretches in strong linkage disequilibrium.

We can analytically calculate the evolutionary dynamics of a population on this fitness landscape in the weak mutation limit, *Nµ* ≪ 1, where *N* is the effective population size and *µ* is the mutation rate per genome per generation. In this limit, the population is monomorphic and individual mutations either rapidly go to fixation or are lost to drift [34]. The dynamics of an evolving population in this limit can be mathematically represented as a Markov process, where the state of the Markov process at time *t* corresponds to the predominant genotype present in the population at that time, and a transition to a new state corresponds to the fixation of a new mutation [34, 35]. For sufficiently large times *t*, the Markov process reaches stationarity, at which point its probability to reside in any given state is provided by the equilibrium distribution *p*_*k*_. We can interpret *p*_*k*_ as the probability to observe the population centered around a genotype carrying *k* mutations at any point in time.

Assuming that we know the equilibrium distribution *p*_*k*_, we can calculate the population mean fitness *f*_eq_ by averaging over the stationary distribution of *k*-mutants,

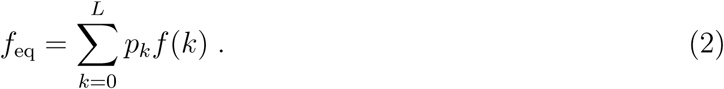

Importantly, *f*_eq_ represents an average over time. It is the mean fitness in the population when averaged both over all individuals in the population and over a long period of time.

The distribution *p*_*k*_ can be calculated using the transition probability *P* (0 → *k*) in the Markov process, solving the detailed balance equations [35]. More precisely, detailed balance entails that in a process where the transitions *i* → *j* and *j* → *i* are both possible, the number of changes *n*_*i*→*j*_ must equal the number *n*_*j*→*i*_. To obtain *n*_*i*→*j*_ and *n*_*i*→*j*_, we calculate the number of *k*-mutants that go to fixation, starting with the wild type sequence with fitness *f*(0), and compare this with the number of *k*-mutants that are replaced by the wild type. Using the Sella-Hirsh fixation formula [35] that is appropriate for a haploid Wright-Fisher process, for a *k*-mutant with fitness *f*(*k*), we find

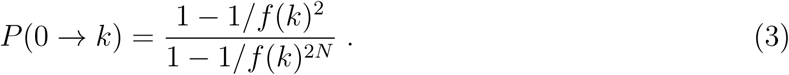

The reverse rate is then

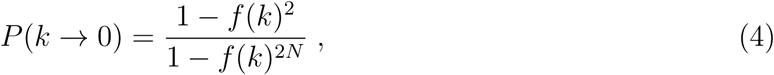

so that the detailed balance condition becomes

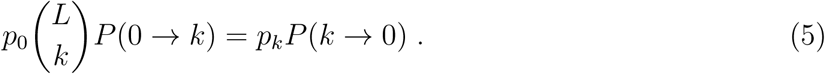

In Eq. (5), *p*_0_ is the equilibrium density of the wild-type, while *p*_*k*_ is the (combined) equilibrium density of *all* individual *k*-mutants. Equations (3–5) then lead to the solution [35]

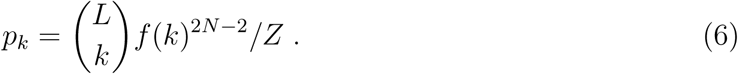

In this expression, *Z* is the partition function

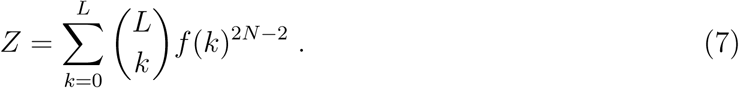

For *q* = 1 we can obtain a closed-form expression for the equilibrium fitness,

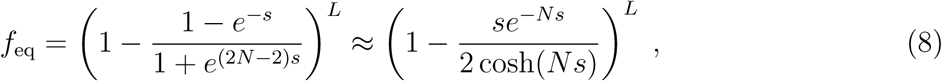

that shows clearly the steep fitness drop with decreasing population size that is due to genetic drift. But different values for *q* affect the fitness drop differently. In Fig. 1A, we can see the dependence of *f*_eq_ on the population size for the multiplicative model (*q* = 1), a model with positive epistasis (*q* = 2.0), as well as the case of negative epistasis (*q* = 0.5), evaluated at *s* = 0.01 and *L* = 100. The model suggests that while positive epistasis protects from a fitness drop for moderate population sizes (higher mean equilibrium fitness), the drop becomes severe once populations dwindle below 100. In fact, plotting *f*_eq_ against *q* as in Fig. 1B reveals a fitness *minimum* as a function of *q*, suggesting that fitness loss via drift can be prevented in two different ways: high positive epistasis or high negative epistasis, while populations with weak or n the critical epistasis parameter o epistasis appear to be the most vulnerable.

**Figure 1:**
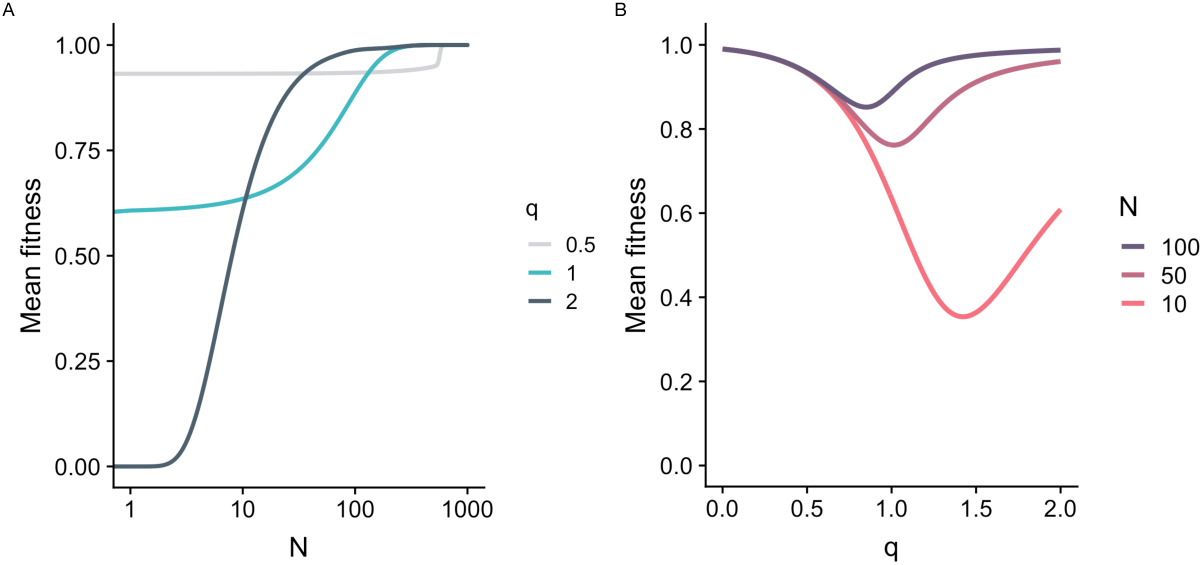
(A) Equilibrium fitness Eq. (2) as a function of population size *N* for strong positive epistasis (*q* = 2.0, dark grey), no epistasis (*q* = 1.0, teal), and strong negative epistasis (*q* = 0.5, light grey). (B) Equilibrium fitness as a function of epistasis *q*, for three different population sizes (red: *N* = 10, dark red: *N* = 50, black: *N* = 100). In both panels, we used *s* = 0.01 and *L* = 100. The equilibrium fitness tends to increase with increasing *N* but displays a minimum at intermediate *q*.

### Two regimes: selection and neutral drift

The minimum in mean equilibrium fitness apparent in Fig. 1B can be seen as interpolating between two regimes: the *neutral drift* regime and the *selection* regime. To formalize these two regimes, we define the critical epistasis parameter *q*^⋆^ at which mean fitness is minimal. Then, the neutral drift regime corresponds to *q* ≪ *q*^⋆^ and the selection regime corresponds to *q* ≫ *q*^⋆^. In the selection regime, an organism’s fitness declines rapidly with increasing number of mutations, and this rapid decline effectively limits the maximum number of mutations an organism can carry. By contrast, in the neutral drift regime additional mutations have increasingly smaller effects on organism fitness, and as a consequence selection cannot effectively purge deleterious mutations.

When *q* ≪ *q*^⋆^, selection cannot effectively purge deleterious mutations, and consequently the evolutionary dynamics are dominated by neutral drift. We can estimate the mean equilibrium fitness in the neutral regime by using

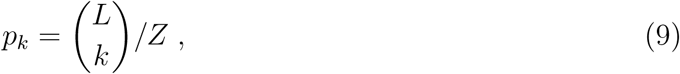

that is, the distribution given by Eq. (6) but with *f*(*k*) ≡ 1. To calculate the mean fitness under this distribution, we insert this expression for *p*_*k*_ into Eq. (2), but note that we need to keep the original expression for *f*(*k*) in Eq. (2). The idea is that in the drift regime fitness differences are sufficiently small that they have no influence on the mutant distribution *p*_*k*_. This does not mean, however, that all organisms have a fitness of 1. The result of this derivation is the dashed line in Fig. 2, which agrees with the full model for sufficiently small *q*.

**Figure 2:**
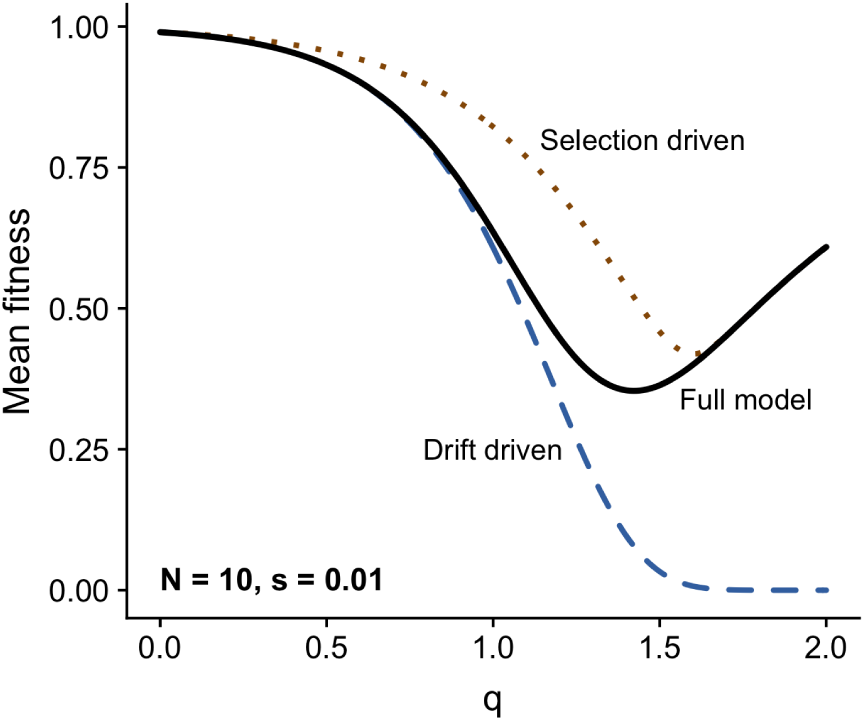
Equilibrium fitness as a function of the epistasis parameter *q* for *N* = 10, *s* = 0.01, and *L* = 100. The solid black line represents the full model, Eq. (2), while the dashed blue line represents the drift-driven approximation, Eq. (9), and the dotted brown line represents the selection-driven approximation, Eq. (10) with a maximum number of mutations *k*_max_ = 20. For *q* ≲ 0.8, the solid black line lies exactly on top of the dashed blue line, and for *q* ≳ 1.7, the solid black line lies exactly on top of the dotted brown line. Thus, the two approximations represent the full model well in the two limits of small and large *q*, respectively.

On the other hand, when epistasis between deleterious mutations is synergistic (*q* ≫ *q*^⋆^), the number of mutations that a population can sustain before it goes extinct is limited to some number *k*_max_. We can model this limitation by imposing a maximum number of mutations *k*_max_,

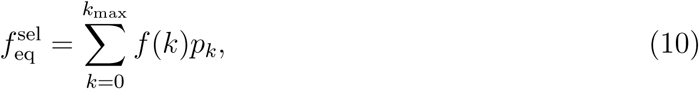

where *k*_max_ *< L*. This truncated mutation model agrees with the full solution for large *q* (Fig. 2, dotted line).

One of the most striking features of the interplay between the neutral regime and the selection regime is the appearance of a minimum mean fitness (as a function of epistasis) where the drop of fitness is largest. The location of this minimum *q*^⋆^ (reflecting the amount of directional epistasis that leads to the largest fitness loss) depends on the population size, the mean deleterious effect of mutations, and the number of loci (Fig. 3).

**Figure 3:**
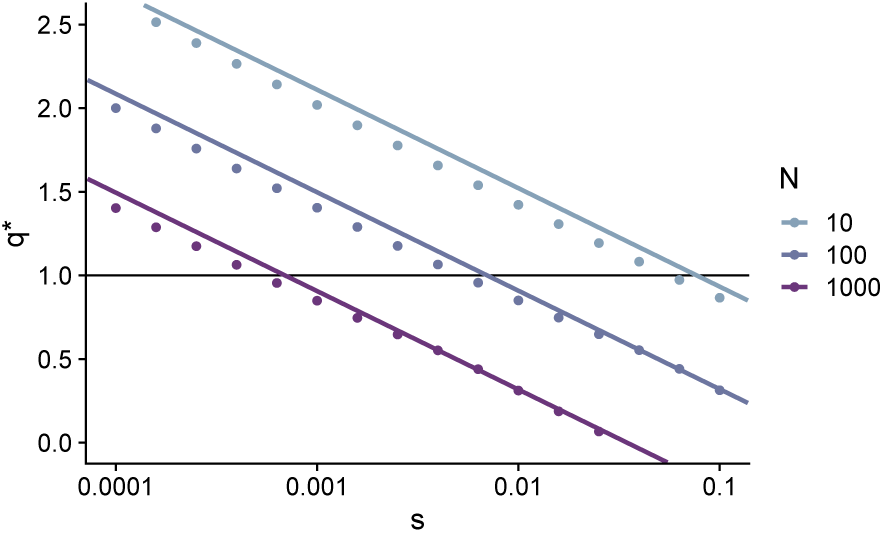
Relationship between the critical epistasis value *q*^⋆^ and the selection coefficient *s*, for different population sizes *N* (*L* = 100 throughout). The lines represent analytically derived *q*^⋆^, as given by Eq. (12). The dots represent *q*^⋆^ values obtained by numerically minimizing Eq. (2). Throughout the entire parameter range, Eq. (12) provides a good approximation to the true location of the minimum.

To estimate the epistasis coefficient at which the steady-state fitness is at its minimum, we analyze the stationary distribution of fitness, Eq. (6), which apart from the normalization constant *Z* consists of two factors, 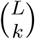 and *f*(*k*)^2*N−*2^ = exp[−*sk*^*q*^(2*N* − 2)]. As discussed in the derivations to Eqs. 9 and 10, these two factors represent neutral drift and selection, respectively. Importantly, for most values of *k* the binomial coefficient 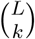 is much larger than 1 whereas the selection term exp[−*sk*^*q*^(2*N* − 2)] is much smaller than 1. Further, to the left of the minimum the binomial coefficient dominates the product whereas to the right of the minimum the selection term dominates. Thus, at the minimum we expect the two factors to cancel, i.e., have a co m bined value of ∼ 1. To arrive at an expression that is independent of *k*, we maximize 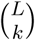 by setting *k* = *L/*2. Then, the condition for maximal fitness loss (where drift maximally balances selection) becomes

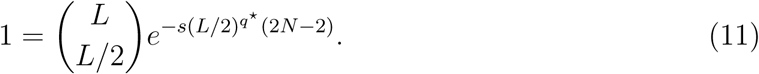

We can solve this equation for *q*^⋆^ by using the Stirling approximation (log *n*! = *n* log *n* − *n*) to expand the binomial coefficient. We obtain for the minimum *q*^⋆^ that

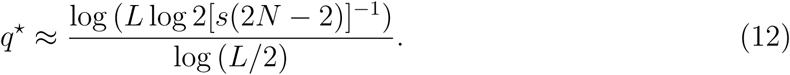

We test this estimate by comparing it to the numerically inferred minimum obtained via numerically minimizing Eq. (2) and find that Eq. (12) generally performs well, though it has a tendency to overestimate the true value of *q*^⋆^ by a few percent (Fig. 3). Importantly, Eq. (12) captures the correct functional relationship between *q*^⋆^ and the model parameters. In particular, the location of *q*^⋆^ is primarily determined by the product of *s* and *N*, and not by their individual values. Further, because the product *sN* enters the expression for *q*^⋆^ via a double-log, *q*^⋆^ changes very slowly even if *sN* changes by orders of magnitude.

### Increased mutation rate exacerbates fitness loss in neutral regime

The theoretical results shown above were derived in the weak mutation limit where every mutation is either lost or goes to fixation before another mutation occurs in the population. In this section we study how finite mutation rates modify those results.

We simulate finite populations on a single-peak fitness landscape at finite mutation rates *µ* using stochastic simulation. The population evolves asexually, and the population size is held constant over time for all simulations. For each combination of mutation rates and epistasis parameters, we simulated populations sizes *N* = 10 and *N* = 100, as well as selection coefficients *s* = 0.01 and *s* = 0.001. We recorded the mean fitness of the population over a period of time after a population reached steady-state (see Methods), as a proxy for this equilibrium fitness *f*_eq_. The simulations of the evolutionary process on the fitness landscape defined by Eq. (1) recover the theoretical results well for small mutation rates, as expected. As the mutation rate increases, we see notable departures from the weak mutation limit for the selection regime (larger *q*), while the neutral (drift) regime is largely unaffected by the increased rates (Fig. 4).

**Figure 4:**
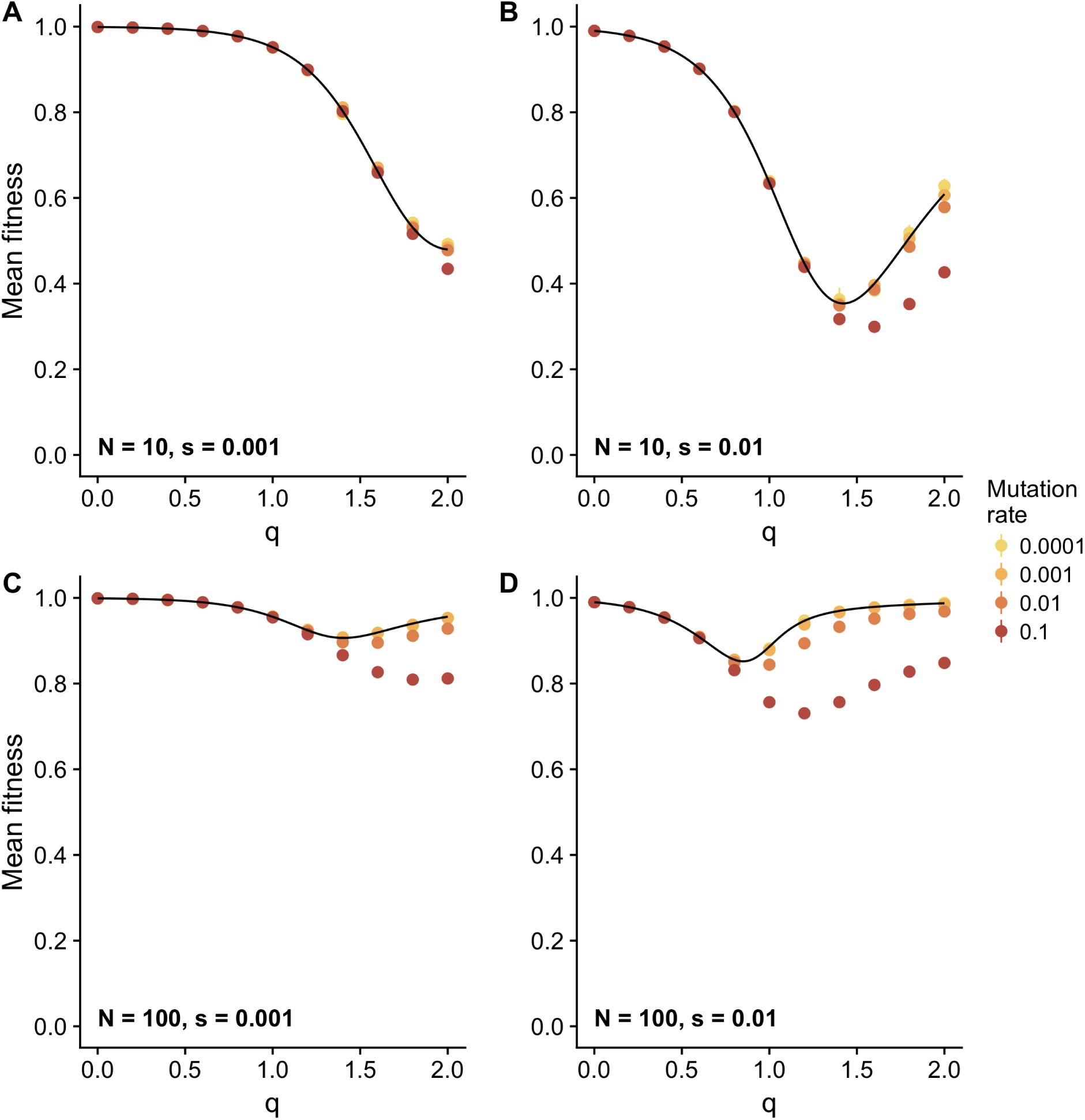
Comparison between theoretical equilibrium fitness and simulated equilibrium fitness. The black line represents theoretical fitness of the population at steady state, Eq. (2). The dots represent mean fitness of a population at steady state over 10 simulations for different mutation rates (see legend). The error bars represent the standard error. For almost all cases, error bars are smaller than the symbol size. (A–D) Theoretical fitness and simulated fitness for different population sizes (*N*), epistasis coefficient (*q*), and selection coefficient (*s*). Mutation rate seems to have no effect on equilibrium fitness to the left of the fitness minimum, but to the right higher mutation rates lead to systematically lower equilibrium fitness values.

In particular, we notice that the minimum of the equilibrium fitness shifts towards higher *q* (Fig. 4). Furthermore, while for small mutation rates an increased epistasis protects from the loss of fitness due to genetic drift (mean fitness does not drop appreciably), it is clear that higher mutation rates negate this effect, and instead exacerbate the loss of fitness. Indeed, the increased mutation rate mimics the effect of a smaller population size (see Fig. 1B), which is expected as the effective population size decreases with mutation rate.

While the depressed equilibrium fitness suggests that there are two routes to withstand genetic drift at small population sizes, it is not clear whether evolutionary trajectories could indeed bifurcate.

### Bifurcation analysis of survival strategies

The minimum in *f*_eq_ at *q*^⋆^ suggests that if *q* were a dynamical variable, then *q*^⋆^ represents an unstable fixed point of the evolutionary dynamics. While *q* is not a dynamical variable in the usual sense, we can simulate it by endowing each genotype with a particular value of *q* that can be changed via mutation. In such a simulation, the statistics of the mutational process affecting *q* (the rate of change *µ*_*q*_ as well as the mean change per mutation Δ*q*) matter, so we test multiple different values for each.

It is worth pointing out that a genotype-dependent *q* appears to contradict the idea of fitness optimization in a landscape with a fixed fitness function such as Eq. (1). Such a function suggests that as a population climbs this peak, the parameters *q* and *s* are unaffected by this climb. While this is true for such a simple fitness function, it does not hold for more realistic evolutionary landscapes (for example in digital life [16,36–38]), where the mean effect of mutations *s* and the directional epistasis *q* are not fixed properties of the landscape, but instead emerge as properties of the local neighborhood in genetic space. As a consequence, moving in this space (via mutations) will affect both *s* and *q*. We attempt to simulate part of that dynamics by allowing *q* to adapt (while keeping *s* fixed). If selection favors a particular value of epistasis, we should see a gradual change in the mean epistasis 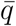 of a population.

In Fig. 5, we show how the mean epistasis parameter 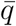 (averaged over sequences in the population) changes over time when populations are seeded with different seed organisms with fixed initial *q*. A bifurcation is indicated when trajectories move to different future fixed points given different initial states. While we can see clear signs of a bifurcation when plotting the mean trajectory in *q*-space over time (Fig. 5), viewing each trajectory separately reveals significant variation among them. In particular, for trajectories that are initialized with a *q* above the fixed point, some trajectories still move towards the low-*q* fixed point, which results in the mean of trajectories to appear constant (for example, in Fig. 5C). We discuss this phenomenon below.

**Figure 5:**
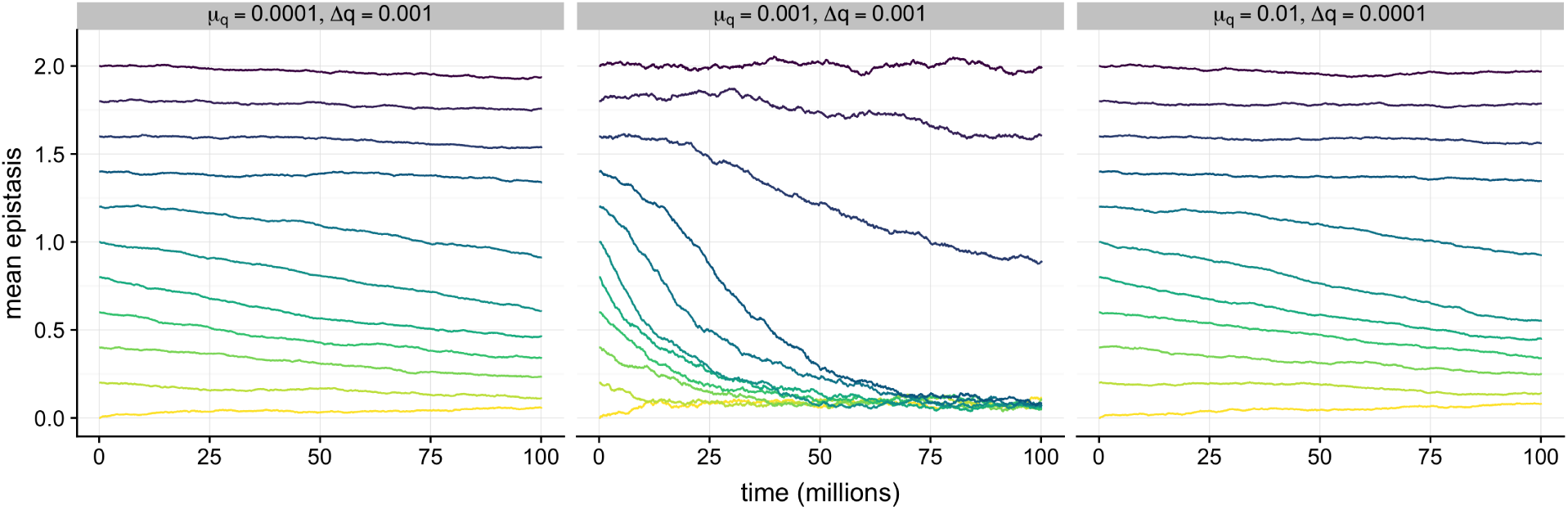
Mean epistasis *q* as a function of time *t* for different combinations of *µ*_*q*_ and Δ*q* in simulations with evolving *q*. Each line corresponds to the mean over 10 replicates. The labels on top of each panel specify the mutation rate of *q, µ*_*q*_, and the mutation step size, Δ*q*. The different colors correspond to different starting points of q. Each population had a fixed *q* until *t* = 200, 000, for equilibration, and then *q* was allowed to evolve. Other simulation parameters were *µ* = 0.01, *N* = 100, *s* = 0.01, and *L* = 100. We observe bistable behavior, such that populations with a mean *q* below the critical value experience continued decline in mean *q*, whereas populations with a mean *q* above the critical value do not.

## Discussion

The dynamics of evolution in asexual population is well-understood in the common population-genetic limits, namely vanishingly small mutation rate and large population (weak mutation and strong selection). When mutation rates are high and selection is weak, the classic theoretical results are undermined by new effects such as mutational robustness (effect of large mutation rate) and drift robustness (effect of small population size), as anticipated by generalized population-genetic models such as “free-fitness” evolution [35, 39–41]. Such theoretical models posit that Darwinian evolution does not optimize reproduction rate, but rather a combination of terms (the “free fitness”, in analogy to the free energy concept of statistical physics) that includes the reproduction rate as well as a term proportional to the inverse of population size and one proportional to mutation rate. In such theories, it is possible to increase the free fitness by trading reproduction rate for robustness to mutations, to drift, or both.

In most populations, we expect both mutational and drift robustness to contribute to survival. For example, when the mutation rate is large, the effective population size is diminished, so that both mutational and drift robustness are bound to be intertwined. The mean directional epistasis between mutations plays a role in both effects. While fitness peaks for mutationally robust populations tend to be flatter with little epistasis between mutations, we also observe something akin to truncation selection [9,10]. In drift robustness, we observe both an increase in neutral mutations as well as an increase in strongly deleterious and lethal mutations, mediated by strong negative epistasis (*q* > 1).

Here, we have calculated the mean equilibrium fitness of a population in the limit of small mutation rates using a simple fitness function with variable epistasis and tunable mutation effect-size, and we have found a minimum as a function of the mean directional epistasis parameter *q* that depends on population size. Stochastic simulations of adaptation on this landscape suggest that the minimum also depends on mutation rate. The model further suggests that there are two attractive fixed points for evolutionary dynamics, namely small *q* where mutations become nearly neutral, and large *q* where deleterious mutations interact synergistically. The low-*q* fixed point^2^ can be identified with mutational robustness (*q* ≈ 0). In contrast, the *q* > 1 fixed point is reminiscent of drift robustness.

While the existence of a minimum in equilibrium fitness is suggestive of an unstable fixed point *q*^⋆^ at which evolutionary trajectories bifurcate towards a low-*q* and a high-*q* fixed point, an agent-based simulation of such trajectories in a bit-string fitness model implementing Eq. (1) but with variable *q* paints a more complicated picture. It is clear from inspection of Eq. (1) that for every single sequence, a reduction of *q* while keeping *k* constant increases fitness (as *∂f/∂q* < 0), no matter what the mean *q* of the population. This means that the population will sense an evolutionary pressure to reduce *q* independently of the mean population epistasis. However, if *q* > *q*^⋆^, a secondary selective pressure appears that acts via the fitness distribution of a sequence’s offspring. For sequences with *q* > *q*^⋆^, sequences with higher *q* have on average off-spring with higher fitness than those with lower *q*, leading to a second-order selective pressure to increase *q*. However, in any particular fitness trajectory, there is a chance that a sequence with *q* < *q*^⋆^ is among the offspring. Such a sequence may then go to fixation and abrogate the evolutionary trajectories leading towards a *q* > *q*^⋆^, even though the selective pressure towards higher *q* is still present. We also expect that the likelihood of mutations that create sequences with *q* < *q*^⋆^ in the offspring distribution depends on *µ*_*q*_ as well as Δ*q*. This is precisely what we observe in Figs. 5 and 6: while for small *q* < *q*^⋆^ the trend towards the mutationally robust fixed point is evident, at *q* > *q*^⋆^ the mean epistasis across 10 replicate experiments often shows a decrease (or remains constant) even though theoretically we expect an approach towards the drift-robust fixed point. The distribution of fitness trajectories shown in Fig. 6 shows that while some trajectories indeed move towards higher *q*, the possibility of mutating towards *q* < *q*^⋆^ leads to trajectories in which the secondary selective pressure towards higher *q* is muted. Indeed, trajectories towards *q* > *q*^⋆^ are absent among the replicates with initial *q* < *q*^⋆^, reinforcing the conclusion that a critical amount of epistasis separates a population’s response to evolutionary stress either in a mutationally-robust, or a drift-robust manner.

**Figure 6:**
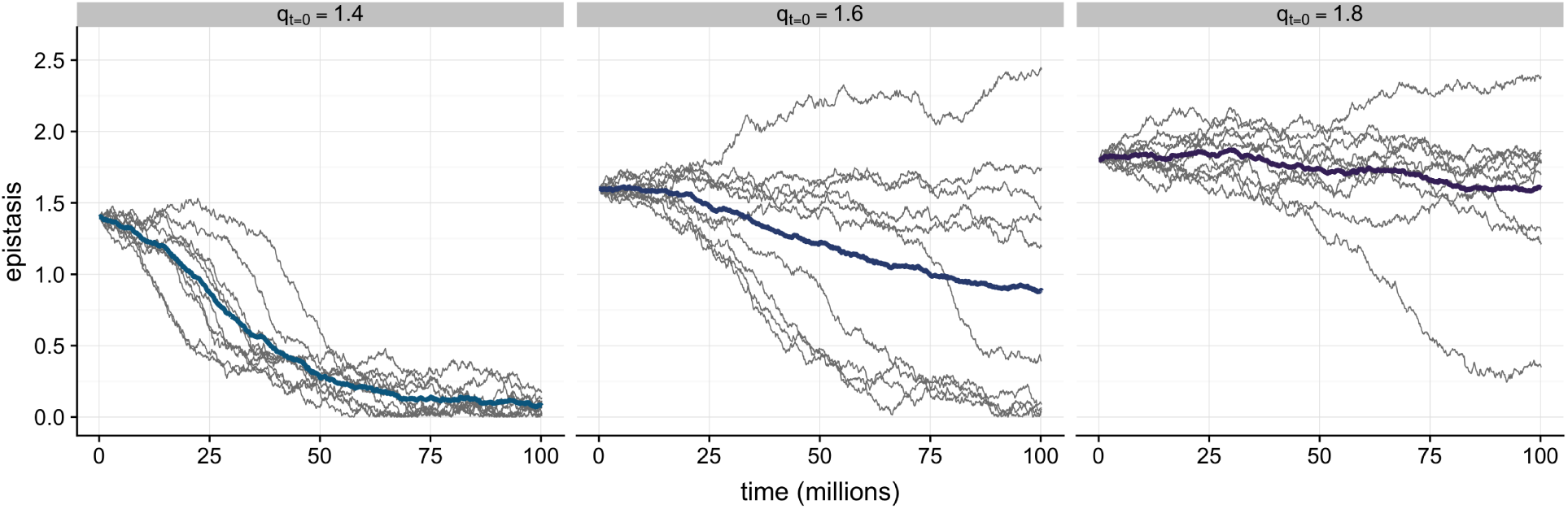
Individual population trajectories for the simulations shown in the middle panel of Fig. 5 (*µ*_*q*_ = 0.001, Δ*q* = 0.001). Thin gray lines correspond to the evolution of individual populations, and the thick colored lines trace the mean among the replicate populations, as in Fig. 5. The labels on top of each panel indicate the initial population epistasis *q*(*t* = 0) in that panel.

Throughout this work, we have used relatively small population sizes around *N* = 100 or less, as any investigation of drift robustness necessitates sufficiently small populations sizes such that drift can play a significant role in the evolutionary dynamics. However, our analytical results provide a more nuanced picture of the conditions under which our results may be relevant. First, we can see from Eq. (12) that the location of *q*^⋆^ depends only on the product *sN* (i.e., the scaled selection coefficient) for sufficiently large *N*. Thus, drift robustness can act even for very large population sizes as long as *s* is sufficiently small. However, this is only true with one additional caveat: There can only be a fitness minimum separating the drift and selection regimes if the number of sites *L* is sufficiently large (on the order of 1*/s*), so that a large number of deleterious mutations can accumulate. By contrast, if both *s* and *L* are small, then the mean fitness is always approximately 1 and whether mutations are present or absent in a genotype makes virtually no difference.

While it is difficult to extrapolate results obtained using the abstract fitness function Eq. (1) to more complex landscapes in which many different peaks with different effect sizes and directional epistasis exist at the same time, our results support the notion that mutational robustness and drift robustness are indeed two different effects, which are likely to be intertwined in realistic scenarios. In particular, it would be interesting to study the response of experimental populations exposed to different mutation rates and populations, something that is possible using strains of *T. brucei*, for example. In those eukaryotic parasites, the directional epistasis between mutations in mitochondrial genes is controlled in part by RNA editing leading to overlapping genes [42]. Because the rate of gene overlap strongly correlates with directional epistasis, the present theory predicts that strains that differ in the number of overlapping genes could take different evolutionary trajectories when subjected to severe bottlenecks. While experimental evolution over prolonged time with parasites through controlled bottlenecks is difficult, such experiments might reveal to us these hidden dimensions of genomic adaptation.

## Methods

### Evolutionary model with fixed epistasis

For all simulations, we implemented an individual-based bit string model. Each individual was represented with a number of deleterious mutations it possesses. The number of deleterious mutations is *k* and *k* = 0, 1, …, *L. L* is the maximum number of deleterious mutations allowed in a population. A population was represented as a vector *V* of a length *L* + 1. Each bin *V*_*k*_ within a vector corresponded to the number of individuals with deleterious mutations *k*. The fitness of an individual could be determined with equation 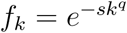, where *s* is the selection coefficient, and *q* is the epistasis coefficient. At each time point, a population reproduces and mutates. A reproduction event increases or decreases the number of individuals within each bin. The number of offspring was drawn from a multivariate distribution, and the probability of reproduction for each bin was determined by the number of individuals within a bin and a bin’s fitness. For all reproduction events, the population size was held constant. After reproduction, a mutation event moves individuals up or down a bin. For most simulations, the maximum number of moves up or down a bin was set to 3. When mutation rate was set to 1, the maximum number of moves was set to 4. The probability of mutating was calculated with

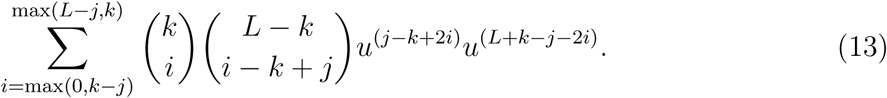

Here, *u* is the per-site mutation rate, *k* is number of mutation an individual has at time *t*, and *j* is the number of mutations an individual has at time *t* + 1. Using the probabilities calculated from the equation above, the number of individuals that would move or stay were drawn from a multivariate distribution. We simulated population sizes of 100 and 10. We set selection coefficients to 0.01 and 0.001. We use mutation rates *µ* of 0.1, 0.01, 0.001, and 0.0001 (mutation rate is defined as the expected number of mutations per genome per duplication, *µ* = *uL*). For each combination of population size, selection coefficient, and mutation rate, we simulated 10 replicates. After a population has reached an equilibrium at *t* = 1, 500, 000, we calculated equilibrium fitness by taking the mean of population fitness over the next 1,000,000 time steps.

### Evolutionary model with evolving epistasis

Similarly to simulations with fixed epistasis, we implemented an individual-based bit-string model to simulate populations with evolving epistasis. A population was represented with two vectors. The length of each of the vectors corresponded to the size of the population, and each element within a vector represented an individual. The first vector contained a number of mutations (*k*) an individual possesses, and the second vector contained an epistasis coefficient (*q*) an individual’s genome experiences. The fitness of an individual could be determined with equation 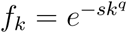. At each time point, a population reproduced and mutated. We reproduced a population by creating a new generation with Wright-Fisher model. We mutated a population with a two step process. First, an individual either gained a mutation (*k* + 1), lost a mutation (*k* − 1), or did not mutate. The probability of mutating was calculated with equation 13. Second, each individual’s epistasis mutated. If epistasis was set to change, it could equally likely increase or decrease by a fixed amount (Δ*q*). We set Δ*q* to remain the same across generations per one trajectory of an evolving population. However, epistasis was only set to evolve after a population has reached an equilibrium (after *t* = 200, 000).

## Data analysis and code

We wrote our simulations in Python [43], using the NumPy [44] and SymPy [45] libraries for numeric and symbolic manipulations of matrices, respectively. Downstream data analysis and visualization was performed in R [46], making extensive use of the tidyverse family of packages [47]. Our simulation and analysis code is available at https://github.com/clauswilke/epistasis_evolution/ and it is archived on Zenodo at https://doi.org/10.5281/zenodo.3558802. Simulation datasets generated with this code are archived in the Texas Data Repository at https://doi.org/10.18738/T8/GUNX76.

## Acknowledgements

This work was supported in part by the National Science Foundation’s BEACON Center for the Study of Evolution in Action, under contract No. DBI-0939454.

## Footnotes

1 The present model in which fitness declines as a function of genetic distance from the wild-type (modulated by epistasis) gives rise to conclusions similar to what Fisher’s geometric model would predict, even though in Fisher’s model the distance from wild-type is phenotypic rather than genetic [32].

2 Note that while technically the low-*q* fixed point is *q* = 0, this value cannot be attained in any realistic population as such a landscape is completely neutral (*f* = 1) in this limit.

